# ECT2 associated to PRICKLE1 are poor-prognosis markers in triple-negative breast cancer

**DOI:** 10.1101/496026

**Authors:** Avais M. Daulat, Pascal Finetti, Diego Revinski, Mônica Silveira Wagner, Luc Camoin, Stéphane Audebert, Daniel Birnbaum, Laurent Kodjabachian, Jean-Paul Borg, François Bertucci

**Affiliations:** Centre de Recherche en Cancérologie de Marseille, Aix Marseille Univ UM105, Inst Paoli Calmettes, UMR7258 CNRS, U1068 INSERM, «Cell Polarity, Cell signalling and Cancer - Equipe labellisée Ligue Contre le Cancer », Marseille, France; Centre de Recherche en Cancérologie de Marseille, Aix Marseille Univ UM105, Inst Paoli Calmettes, UMR7258 CNRS, U1068 INSERM, «Predictive Oncology team », Marseille, France; Aix Marseille University, CNRS, INSERM, Institut Paoli-Calmettes, CRCM, Marseille Protéomique, Marseille, France; Aix Marseille Univ, CNRS, IBDM, Marseille, France

**Keywords:** PRICKLE1, ECT2, signaling, therapeutic targets, Triple Negative Breast Cancer

## Abstract

**Background:** Triple-negative breast cancers are poor-prognosis tumors characterized by absence of molecular signature and are chemotherapy is still the only systemic treatment. Currently, research focus to identify biomarkers that may be usable for prognosis and/or for treatment, notably among the proteins involved in cell migration and metastatic capacity.

**Methods:** We used proteomic approach to identify protein complexes associated to PRICKLE1 and the mRNA expression level of the corresponding genes in a retrospective series of 8,982 clinically annotated patients with invasive primary breast cancer were assessed. Then, we characterize molecularly the interaction between PRICKLE1 and the guanine nucleotide exchange factor ECT2. Finally, experiments in Xenopus have been carrying out to determine their evolutionary conserved interaction.

**Results:** We have identified a network of proteins interacting with the prometastatic scaffold protein PRICKLE1 that includes several small G-protein regulators involved in cell migration and metastasis. Combined analysis expression of PRICKLE1 and small G-protein regulators expression has a strong prognostic value in TNBC. We show that PRICKLE1 controls the activity of ECT2 on RAC1 signaling, a pathway required for cancer cell dissemination.

**Conclusions:** This work supports the idea that promigratory proteins, which are overexpressed in cancerous epithelium, are suitable pharmaceutical targets.

## Introduction

Triple-negative breast cancer (TNBC) is the most aggressive molecular subtype of breast cancer^1^. In contrast with mammary cancers of other subtypes (HR+/HER2− and HER2+), TNBCs do not express hormone receptors and HER2 oncogene and thus are not candidate to hormone therapy and anti-HER2 therapy^1^. Chemotherapy is the only systemic therapy currently approved for this subtype. However, TNBC is highly invasive with strong metastatic propensity^1^. We recently identified *PRICKLE1* as poor-prognosis marker in breast cancer^2^. PRICKLE1 is a member of a conserved group of proteins involved in planar cell polarity (PCP) pathway^3^. This pathway is well characterized in epithelial tissue morphogenesis during embryonic development of invertebrates and vertebrates. The organization of PCP relies on the spatial distribution of proteins at the plasma membrane such as Wnts, Frizzled, Vang Gogh, Flamingo, Dishevelled, Diego, and Prickle. In vertebrates, homologous genes are involved in the regulation of convergent-extension during the early stages of gastrulation which leads to the organization of cells to organize the head-to-tail axis^3, 4^. Prickle1 plays a pivotal role to regulate PCP in Drosophila^5^, as well as convergent-extension in Zebrafish^6^ and Xenopus^7^. PRICKLE1 is an evolutionary conserved cytoplasmic protein and contains from the amino-terminal end a PET followed by three LIM domains and a C-terminal farnesylation site^8^. Recently, we and others have demonstrated the prominent role of PRICKLE1 during cancer progression^2, 9–11^. PRICKLE1 is a prometastatic molecule and regulates oriented cell migration in various cell lines including the MDA-MB-231 prototypal TNBC cell line^2, 10^. At the molecular level, PRICKLE1 regulates subcellular localization of its associated proteins such as VANGL2^8, 12^, RICTOR^2^, ARHGAP22/24^10^, and LL5β^11^ in order to coordinate oriented cellular migration.

Here, we identified the proteome associated to PRICKLE1 in MDA-MB-231 cells. Among the proteins associated to PRICKLE1, our attention was attracted by a large subset of small G-protein regulators. Since regulation of cancer migration depends of the activation of small G-proteins such as Rac, Rho, and Cdc42, we further explored the role of this subset of small G-protein regulators using transcriptomic analysis from publically available data set obtained from patients with breast cancer. We gathered all the identified small G-protein regulators in a metagene to allow an in-depth analysis and showed that *PRICKLE1* was not only overexpressed in TNBC, but its associated proteins were also up-regulated in TNBC and were poor-prognosis markers. To further explore the protein complex associated to PRICKLE1, we focused our attention on the Rho-Guanylyl Exchange Factor (GEF) called Epithelial cell transforming sequence 2 (ECT2). In non-transformed cells, ECT2 regulates cytokinesis by regulating Rac1 activity^13–16^. *ECT2* is frequently up-regulated in various cancers such as ovarian^14^, lung^17^ and breast cancer^18^. Knockdown of ECT2 inhibits Rac1 activity and block transformed growth, invasion and tumorigenicity^13, 16^. Here we showed that PRICKLE1 was associated to ECT2 in MDA-MB-231 cells to regulate Rac1 activity and therefore promote cell motility. Using *Xenopus laevis* embryos, we showed that Prickle1 and Ect2 acted synergistically during embryonic development. Together these data demonstrate the importance of Prickle1 and its associated protein complex as poor-prognosis markers in TNBC and give evidence that PRICKLE1 can be a suitable therapeutic target for treatment of still lacking targeted therapy for this aggressive subtype.

## Materials and Methods

### Rac1 activity assay

Cells were lysed with ice cold lysis buffer (50 mM Tris, pH7.6, 150mM NaCl, 0.1% Triton X-100, 20mM MgCl_2_ supplemented with protease inhibitor (Sigma)). Supernatant were collected after 10 min of centrifucation at 10,000xg at 4°C. Protein concentration is measured from the solubilized fraction and adjusted to 2mg/mL. 10% of the lysates are conserved as loading controls. 100μg of GST-CRIB are added to 2mg of lysate and incubate with rotation during 30 min at 4°C. Beads are then washed with 10 volumes of lysis buffer. Rac-GTP forms are eluted from the beads using 2x Leammli buffer. 30% of the sample are run on 15%SDS-PAGE gel and transfer to PVDF and blot with the indicated antibody.

### Breast cancer samples and gene expression profiling

Our institutional series included 353 tumor samples from pre-treatment invasive primary mammary carcinomas either surgically removed or biopsied.^19^ The study was approved by our institutional review board. Each patient had given a written informed consent for research use. Samples had been profiled using Affymetrix U133 Plus 2.0 human microarrays (Santa Clara, CA, USA). We pooled them with 35 public breast cancer data sets comprising both gene expression profiles generated using DNA microarrays and RNA-Seq and clinicopathological annotations. These sets were collected from the National Center for Biotechnology Information (NCBI)/Genbank GEO, ArrayExpress, European Genome-Phenome Archive, The Cancer Genome Atlas portal (TCGA) databases, and authors’ website (**Supplementary Table 1**). The final pooled data set included 8982 non-redundant non-metastatic, non-inflammatory, primary, invasive breast cancers.

### Gene expression data analysis

Before analysis, several steps of data processing were applied. The first step was the normalization of each set separately. It was done in R using Bioconductor and associated packages; we used quantile normalization for the available processed data from non-Affymetrix-based sets (Agilent, SweGene, and Illumina), and Robust Multichip Average (RMA) with the non-parametric quantile algorithm for the raw data from the Affymetrix-based sets. In the second step, we mapped the hybridization probes across the different technological platforms represented as previously reported.^20^ When multiple probes mapped to the same GeneID, we retained the most variant probe in a particular dataset. We log2-transformed the available TCGA RNA-Seq data that were already normalized. In order to avoid biases related to trans-institutional IHC analyses and thanks to the bimodal distribution of respective mRNA expression levels, the ER, progesterone receptor (PR), and HER2 statutes (negative/positive) were defined on transcriptional data of *ESR1, PGR*, and *HER2* respectively, as previously described.^21^ The molecular subtypes of tumors were defined as HR+/HER2− for ER-positive and/or PR-positive and HER2-negative tumors, HER2+ for HER2-positive tumors, and triple-negative (TN) for ER-negative, PR-negative and HER2-negative tumors. Next, expression levels of *PRICKLE1* and 10 genes of interest from the protein complex associated to Prickle1 (namely, *ARHGAP21, ARGHAP22, ARHGAP23, ARHGEF2, ARHGEF40, BCR, ECT2, IQGAP3, MYO9B*, and *STARD13*) were extracted from each of the 36 normalized data sets. Before analysis, gene expression levels were standardized within each data set using the PAM50 luminal A population as reference. This allowed to exclude biases due to laboratory-specific variations and to population heterogeneity and to make data comparable across all sets. *PRICKLE1* and *ECT2* upregulation in a tumor was defined by an expression level above median expression the other cases being defined as downregulation. GEF/GAP activity was based on metagene approach and computed on the mean of the 10 related genes standardized. GEF/GAP activity “up” was defined by a metagene score value above the global median of the metagene whereas other cases were defined as “down”.

### Statistical analysis

Correlations between tumor classes and clinicopathological variables were analyzed using the one-way analysis of variance (ANOVA) or the Fisher’s exact test when appropriate. Metastasis-free survival (MFS) was calculated from the date of diagnosis until the date of distant relapse. Follow-up was measured from the date of diagnosis to the date of last news for event-free patients. Survivals were calculated using the Kaplan-Meier method and curves were compared with the log-rank test. The likelihood ratio (LR) tests were used to assess the prognostic information provided beyond that of PRICKLE1 model, GEF/GAP metagene or ECT2 model, assuming a x^2^ distribution. Changes in the LR values (LR-ΔX^2^) measured quantitatively the relative amount of information of one model compared with another. All statistical tests were two-sided at the 5% level of significance. Statistical analysis was done using the survival package (version 2.30) in the R software (version 2.15.2; http://www.cran.r-project.org/). We followed the reporting REcommendations for tumor MARKer prognostic studies (REMARK criteria)^22^.

### Xenopus embryo injections, plasmids, RNAs, and Mos

Eggs obtained from NASCO females were fertilized in vitro, dejellied and cultured as described previously^23^. Wild-type embryos were obtained using standard methods^24^ from adult animals and staged according to Nieuwkoop and Faber (1994).

Ect2 riboprobe was generated from *Xenopus laevis* full-length Ect2 cDNA, obtained from Dharmacom™ (Plasmid XGC ect2 cDNA, Clone ID: 5083828; pCMV-SPORT6.ccdb). The cDNA was subcloned in pBS-SK vector. For *Ect2* sense probe the plasmid was linearized by NotI and transcribed with T7 RNA polymerase. For *Ect2* antisense probe the plasmid was linearized by EcoRV and transcribed with T3 RNA polymerase.

Synthetic capped mRFP mRNA was produced using Ambion mMESSAGE mMACHINE Kit. pCS2-mRFP was linearized with NotI and mRNA was synthesized with Sp6 polymerase. 0,5ng of mRFP capped mRNA was used as injection control and tracer.

Morpholino antisense oligonucleotides (MO) were obtained from Genetools^®^, and the sequences were the following: Prickle1 (Pk1) 5’-CCTTCTGATCCATTTCCAAAGGCAT-3’^25^; ECT2 5’-TACTGGGAGAGCCATGTTTGATTT-3’. Embryos at 2-cell stage were injected in each blastomere with various doses of MOs. Embryos were cultured in modified Barth’s solution until stage 28, when they were photographed.

## Results

### Mass spectrometry analysis of the PRICKLE1 complex shows that PRICKLE1 is associated with small G-protein regulators and modulates Rac and Rho activity

We and others have shown that PRICKLE1 contribute to cancer cell dissemination in various cancers^2, 9–11^. To investigate the molecular mechanisms underlying the role of PRICKLE1 in tumorigenicity, and notably cell motility and dissemination, we generated a stable cell line expressing GFP-PRICKLE1 in the MDA-MB-231 highly invasive TNBC cell line. To identify protein complexes associated to PRICKLE1 in these cells, we performed anti-GFP immunoprecipitation followed by mass spectrometry analysis. We identified previously known PRICKLE1 interactors such as VANGL1, MINK1, RICTOR, LL5β, PLK1, and USP9x, validating our approach (**Fig. 1A**). Cell migration is a complex and dynamic process that involves continuous remodeling of the cellular architecture and relies on spatiotemporal modulation of signaling networks including Rho-family GTPases. Our attention was attracted by the large number of regulators of Rho-family GTPases such as Rac1, Rho and Cdc42 (**Fig. 1B**), known to be notably involved in the regulation of cell motility and considered as interesting drug targets to prevent cancer dissemination.

**Figure 1:**
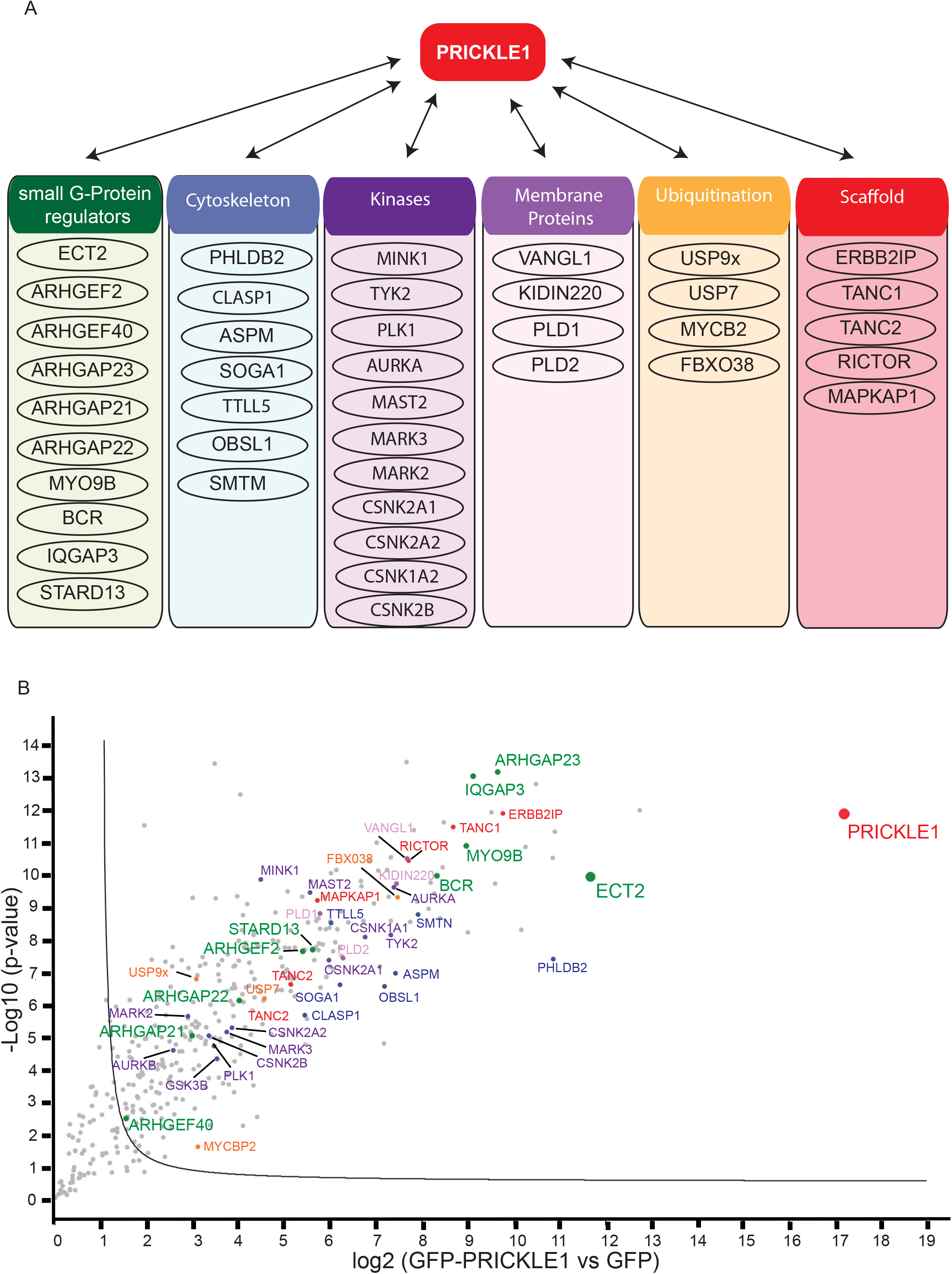
Mass spectrometry analysis of the PRICKLE1 protein complex from a TNBC cell line. A) Schematic representing the proteins associated to PRICKLE1 identified by mass spectrometry analysis from MDA-MB-231 cell extracts. Proteins have been classified following their function in several groups: Small G-proteins regulators, associated to cytoskeleton, Kinases, involved in Ubiquitination process, Membrane integrated, Scaffold proteins and others. B) Volcano plot showing the significance two-sample *t*-test (−Log p-value) *versus* fold-change (Log2 (GFP-PRICKLE1 versus GFP as control)) on the y and x axes, respectively. The full line is indicative of protein hits obtained at a permutation false discovery rate of 1% (pFDR). Data results from two different experiments processed three times. PRICKLE1 (the bait) is represented in red and ECT2 one of the most abundant PRICKLE1 associated partner is represented in green.

### Prognosis value of PRICKLE1-interacting small G-protein regulators in TNBC

Based on our generated proteomic data describing the protein complex associated to PRICKLE1, we focused our attention on the 10 regulators of small G-proteins (i.e. Rho-GEF and Rho-GAP) identified, including ARHGAP21, ARGHAP22, ARHGAP23, ARHGEF2, ARHGEF40, BCR, ECT2, IQGAP3, MYO9B and STARD13. We assessed the mRNA expression level of the corresponding genes in a retrospective series of 8,982 clinically annotated patients with invasive primary breast cancer gathered from several public data bases (**Table S1**). Within these 10 genes, *ECT2, IQGAP3* and *MYO9B* were the most overexpressed in tumors as compared to normal breast (**Fig. 2A**), whereas *ARHGEF40* and *STARD13* showed the lowest expression levels. We built a metagene including these 10 genes and compared its expression level in three molecular subtypes of breast cancer (RH+/HER2−, HER2+, and TN). The metagene was significantly up-regulated in the TN subtype comparatively to the two others subtypes (p<1.0 × 10^−250^, Anova) (**Fig. 2B**).

**Figure 2:**
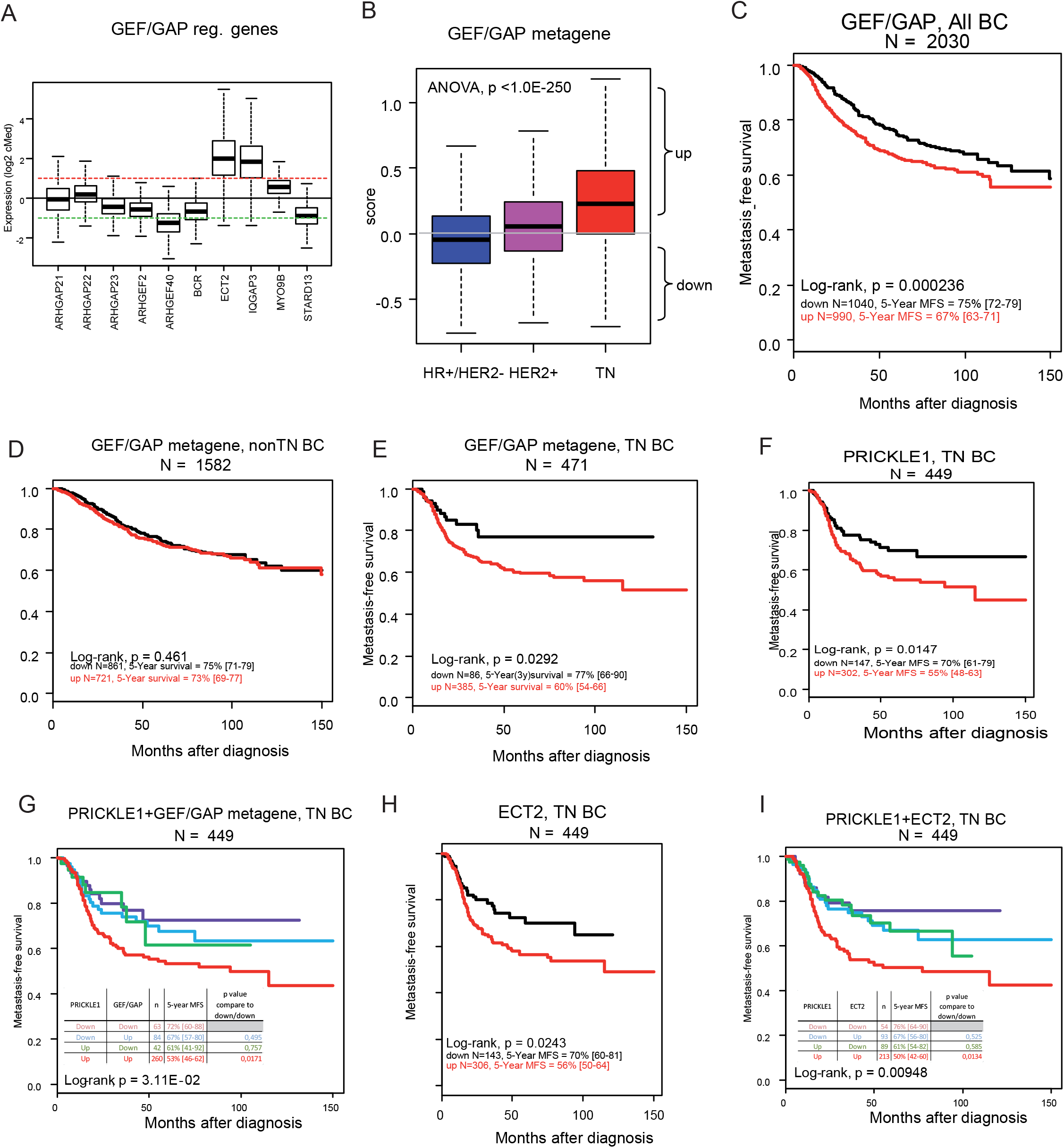
Prognosis value of PRICKLE1-interacting small G-protein regulators in TNBC and cooperation between *PRICKLE1* and *ECT2* as poor prognosis markers. A) Boxplot of GEF/GAP regulators expression across breast cancers. B) Boxplot of GEF/GAP regulators expression across triple negative (TN) *versus* HR+/HER2− or HER2+ breast cancer. C) Kaplan-Meier curves of metastasis-free survival among breast cancers patients according to overexpression (Up) *versus* underexpression (Down) of GEF/GAP metagene mRNA. D) Kaplan-Meier curves of metastasis-free survival among non-TNBC patients for GEF/GAP metagene mRNA expression. E) Kaplan-Meier curves of metastasis-free survival among TNBC patients for GEF/GAP metagene mRNA expression. F) Kaplan-Meier curves of metastasis-free survival among TNBC patients for *PRICKLE1* mRNA expression. G) Kaplan-Meier curves of metastasis-free survival among TNBC patients for *PRICKLE1* mRNA and GEF/GAP metagene expression. H) Kaplan-Meier curves of metastasis-free survival among TNBC patients for *ECT2* mRNA expression. I) Kaplan-Meier curves of metastasis-free survival among TNBC patients for *PRICKLE1* and *ECT2* mRNA expression.

We then searched for correlations between the GAP-GEF metagene expression (as binary variable) and the clinicopathological features of samples, including MFS. Within the 8,982 breast cancer samples analyzed, 4,491 tumors (50%) showed metagene upregulation when compared with normal breast (ratio T/NB ≥ 2; ‘‘metagene-up’’ group), and 4.491 (50%) did not (ratio <2; ‘‘metagene-down’’ group) (**Table 1**). We found significant correlations between the metagene status and patients’ age (p<0.001), grade (p<0.001), ER (p<0.001), PR (p<0.001), and HER2 (p=0.012) statutes and as shown above with molecular subtypes. MFS data were available for 2,030 patients: the 5-year MFS was 75% (95 Cl, 72-79) in the “metagene-down” group *versus* 67% (95Cl, 63-71) in the “metagene-up” group (p=0.00023, log-rank test; **Fig. 2C**). In fact, such prognostic correlation was only observed in TNBC patients, and not in the non-TNBC ones (p=0.461, log rank test; **Fig. 2D**). In TNBC patients, the 5-year MFS was 77% (95 Cl, 66-90) in the “metagene-down” group *versus* 60% (95Cl, 54-66) in the “metagene-up” group (p=0.029, log-rank test; **Fig. 2E**).

**Table 1:**
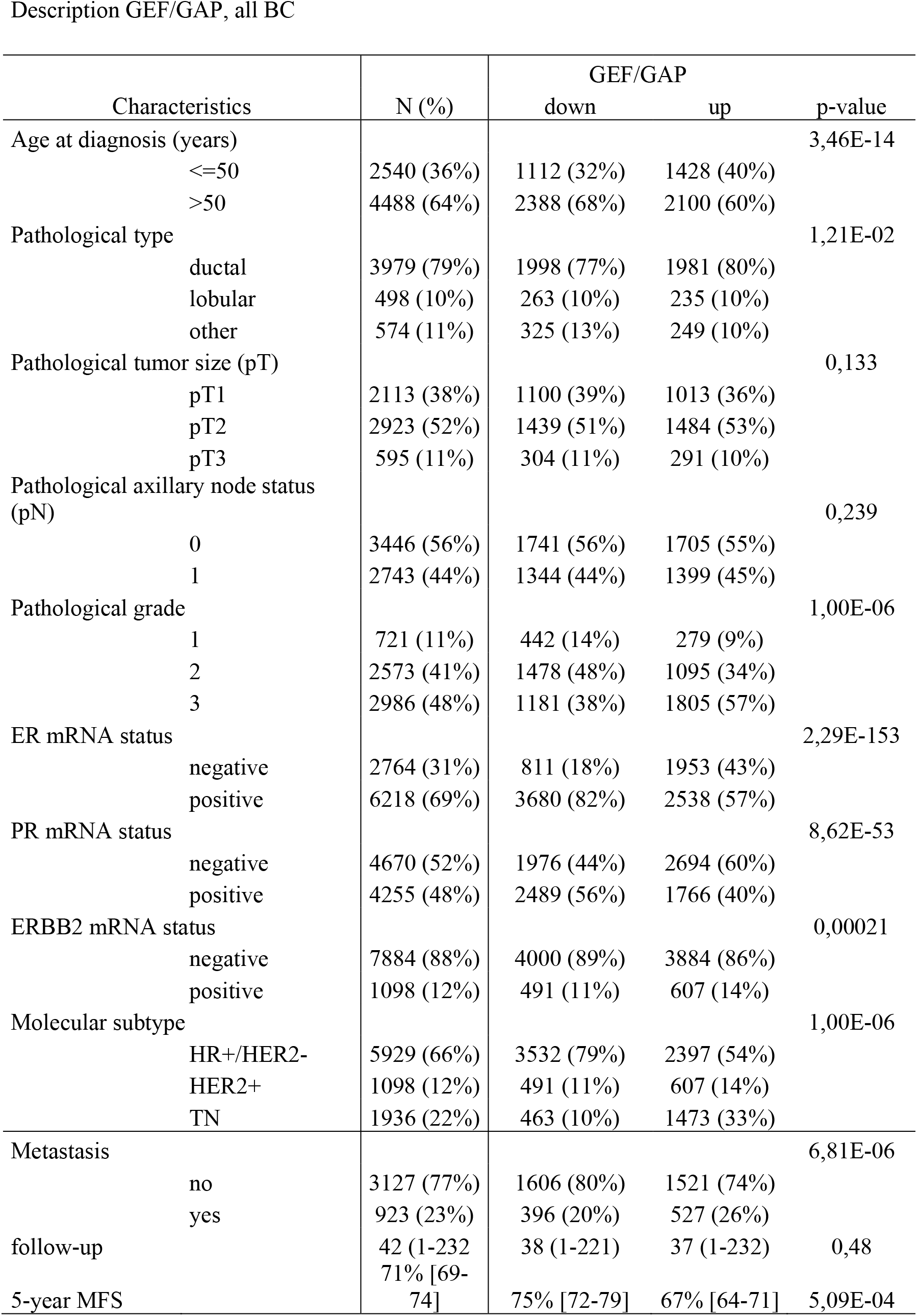
Description GEF/GAP, all BC

### Cooperation between PRICKLE1 and ECT2 as poor-prognosis marker in TNBC

We have previously shown that *PRICKLE1* upregulation is associated with poor MFS in basal breast cancer^2^, a molecular subtype mainly composed of TNBC. In the present series of TNBC, we confirmed that *PRICKLE1* upregulation was associated with shorter metastasis free survival (MFS), with 70% 5-year MFS (95Cl, 61-79) *versus* 55% (95Cl, 48-63) in the *PRICKLE1*-down group and the *PRICKLE1*-up group respectively (p=0.0147, log-rank test) (**Fig. 2F**). Since *PRICKLE1* and the 10 genes of the metagene interact together, we searched for an eventual cooperation of their association in prognostic term. First, we analyzed the combination of the metagene expression and *PRICKLE1* expression. Interestingly, patients with upregulation of both markers displayed shorter 5-year MFS (53%, 95Cl, 46-62) than patients without upregulation of both markers (72%, 95Cl 60-88; p=0.017, log-rank test), whereas patients with intermediate status (up and no-up, and vice-versa) showed intermediate 5-year MFS not significantly different from the same patients (p=0.757, and p=0.495 respectively, log-rank test; **Fig. 2G**). This data suggest that metagene expression and *PRICKLE1* expression might provide complementary prognostic value. Such complementarity between the two prognostic variables was tested in the TN patients using the likelihood ratio (LR) test. As shown in **Table 2A**, the metagene tended to add prognostic information to that provided by *PRICKLE1* expression (LR-ΔX2=2.75, p=0.097).

**Table 2:** Model of comparision

| **<u>MFS,TN BC</u>** | statistic |  | p-value |
| --- | --- | --- | --- |
| PRICKLE1 | LRX<sup>2</sup> | 6,23 | 1,25E-02 |
| PRICKLE1 + GEF/GAP | LRX<sup>2</sup> | 8,98 | 1,12E-02 |
| GEF/GAP+PRICKLE1 vs.<br>PRICKLE1 | ΔLRX<sup>2</sup> | 2,75 | 0,097 |

**B, PRICKLE1 & ECT2**
| **<u>MFS,TN BC</u>** | statistic |  | p-value |
| --- | --- | --- | --- |
| PRICKLE1 | LRX<sup>2</sup> | 6,23 | 1,25E-02 |
| PRICKLE1 + ECT2 | LRX<sup>2</sup> | 11 | 4,14E-03 |
| ECT2+PRICKLE1 vs. PRICKLE1 | ΔLRX<sup>2</sup> | 4,74 | 2,90E-02 |

Second, because ECT2 was one of the most prominent hit identified by mass spectrometry analysis (**Fig. 1B**) and the gene most overexpressed in TNBCs among members of the metagene (**Fig. 2A**), we investigated whether *ECT2* expression alone (without the nine other genes of the metagene) would be sufficient to improve the prognostic value of *PRICKLE1* expression in TNBC patients. As shown in **Figure 2H**, patients with *ECT2* upregulation displayed shorter 5-year MFS (56%, 95Cl 50-64) than patients without upregulation (70%, 95Cl 60-81; p=0.0243, log-rank test). More interestingly, ECT2 expression status increased the prognostic value of *PRICKLE1* expression when combined. Patients with upregulation of both genes displayed 50% 5-year MFS (95Cl, 46-62) *versus* 67% for patients with intermediate status (up and down, and vice-versa) *versus* 76% (95Cl, 64-90) for patients without upregulation of both markers (p=0.0134, log-rank test; **Fig. 2I**). The model comparison (**Table 2B**) showed that such *ECT2* prognostic information added to that of *PRICKLE1* expression was statistically significant (LR-ΔX2=4.74, p=0.029), indicating that *ECT2* expression improved the prognostic value of *PRICKLE1* expression in TNBC.

#### PRICKLE1 binds to ECT2 through it PET domain and modulates Rac1 activity

We then sought to investigate the molecular mechanisms potentially associated to this cooperation of PRICKLE1 and ECT2 expressions to confer poor prognosis. ECT2 is a Rho-GEF and acts in non-cancerous cells as regulator of cytokinesis by exchanging GDP to GTP on the small GTPases, RhoA, Rac1 and Cdc42^26^. ECT2 is upregulated in human cancers and acts as oncogene^27^. In lung and ovarian cancer, ECT2 has a distinct role than cytokinesis and acts in the nucleus by recruiting Rac1 and effectors which are required for tumour initiation and transformation^13, 15, 16^. ECT2 knockdown inhibits Rac1 activity leading to a decrease of tumorigenicity and invasion in lung adenocarcinoma^13^. Recently, ECT2 has been described to be upregulated in breast cancer^18^. At the cellular level, ECT2 is localized in the cytoplasm of cancerous cells^16^. To confirm our mass spectrometry analysis, we immunoprecipitated GFP-PRICKLE1 stably expressed in MDA-MB-231 cells using GFP-targeted antibody and we assessed the presence of ECT2 associated to PRICKLE1 immunoprecipitate by western blot analysis complex (**Fig. 3A**). We confirmed that ECT2 is associated with PRICKLE1 in MDA-MB-231 cells. We further confirmed that ECT2 colocalizes in actin-enriched structures of lamellipodia along with PRICKLE1 using MDA-MB-231 stably expressing GFP-PRICKLE1 (**Fig. 3B**). We next decided to map the domain of interaction between PRICKLE1 and ECT2. We thus generated deleted versions of PRICKLE1 lacking the PET and/or the LIM domains and a construct encompassing the PRICKLE1 C-terminal region. We co-transfected HEK293T cells with the indicated flag tagged PRICKLE1 mutants with Cherry-ECT2. After Flag immunoprecipitation, we assessed the presence of Cherry-ECT2 by western blot analysis. We observed that the PET domain of PRICKLE1 was required for the formation of the PRICKLE1-ECT2 protein complex (**Fig. 3C**).

**Figure 3:**
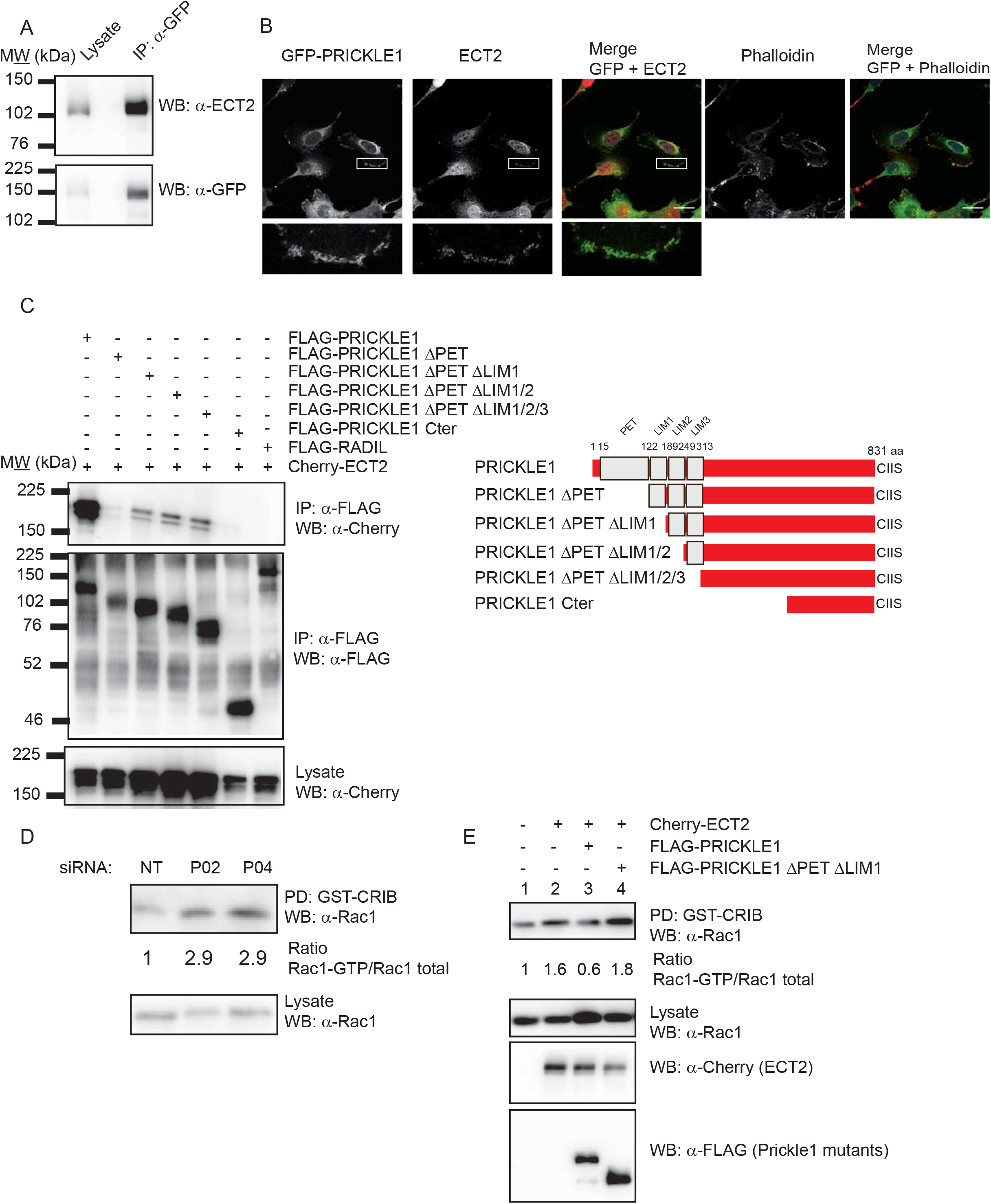
PRICKLE1 is associated to the Rho-GEF ECT2 and controls its activity. A) Immunopurification of GFP-PRICKLE1 from MDA-MB-231 cell lysate using GFP nanobodies coupled to sepharose beads allows the identification of ECT2 associated to PRICKLE1. B) Immunofluorescence of MDA-MB-231 cells stably expressing GFP-PRICKLE1 shows that ECT2 (endogenous) is colocalized with PRICKLE1 and enriched in actin structures within the lamellipodia. C) Mapping of PRICKLE1 domain of interaction with ECT2. HEK293T cells were co-transfected with the indicated form of PRICKLE1 (see on the left for topology details) and Cherry-ECT2. After FLAG immunopurification, presence of ECT2 is detected using anti-cherry antibody. D) Downregulation of PRICKLE1 expression using siRNA targeting *PRICKLE1* shows an increase of Rac activity in MDA-MB-231 cells. E) PRICKLE1 modulates ECT2 activity. Using HEK293T cells, we expressed or co-expressed ECT2 with full length PRICKLE1 or a deleted version of PRICKLE1 lacking its domain of interaction with ECT2. Overexpression of ECT2 leads to an increase of Rac activity which was inhibited when PRICKLE1 is co-expressed. Co-expression of a mutant form of PRICKLE1 did not modify the gain of function observed by ECT2 overexpression.

We further assessed PRICKLE1 contribution on Rac activity. We used previously characterized siRNAs^2^ to specifically downregulate PRICKLE1 expression in MDA-MB-231 cells. We observed that PRICKLE1 modulated Rac1 activity, suggesting a prominent role of PRICKLE1 in the regulation of Rho-GEF and Rho-GAP (**Fig. 3D**). We next set up an essay to monitor the role of PRICKLE1 on ECT2 Rho GEF activity. We expressed cherry-ECT2 in HEK293T cells and observed an increase of active Rac1 (lane 2). However, when flag-PRICKLE1 was co-expressed with cherry-ECT2, we observed an inhibitory effect of PRICKLE1 (lane 3). This observation was confirmed by the co-expression of PRICKLE1 delta PET delta LIM1 which is unable to bind ECT2 and does not affect the gain of activity of ECT2 in our system (lane 4) (**Fig. 3E**). Altogether, our data suggest that PRICKLE1 is associated with ECT2 in actin-rich structures within the lamellipodia of the cells in order to modulate the activity of the ECT2 on Rac1.

#### Prickle1 and Ect2 functionally interact in Xenopus during embryonic development

PRICKLE1 is an evolutionary conserved protein and plays a pivotal role during gastrulation to modulate convergent-extension movements (CE), which are crucial to shape the body plan^7, 28^. To test whether Ect2 is required for the previously characterized function of Prickle1 during CE, we first compared and analyzed the RNA-seq profile of *prickle1* and *ect2* reported on the public XenBase repository^29^ (data not shown). We noticed a sharp peak of zygotic *ect2* expression at stage 9, which decreases abruptly at stage 10, just before gastrulation and CE movements take place. Zygotic *prickle1* expression also begins to increase at stage 9, reaching a maximum at stage 12 (mid gastrula), and gradually decreasing until the end of neurulation. We next performed *in situ* hybridization and detected expression of *ect2* RNA in the animal hemisphere up until stage 9 (**Fig. 4A**). Thus, *ect2* transcription appears to terminate when *prickle1* transcription starts. However, inspection of genome-wide proteomic data^30^ indicated that Ect2 protein levels were maintained during gastrulation, suggesting that Ect2 could cooperate with Prickle1 to regulate morphogenetic movements. To test this hypothesis, we performed Prickle1 and Ect2 knockdown through antisense morpholinos (MO) injections, and assessed CE problems (**Fig. 4B**). Injection of 40ng MO Prickle1 led to CE defects in 73% of embryos, in comparison to non-injected embryos (98%) or embryos injected with RFP as control (83%). This data is consistent with previously published results^7, 12^. We then injected 20ng of MO targeting Ect2 and we observed CE problems at a rate of 71%, phenocopying the effect observed with MO Prickle1 with narrower and shorter embryos at tailbud stage 28. We then defined subthreshold doses of individual Mo-Prickle1 (</=10ng) and Mo-Ect2 (</=10ng) that yielded moderate CE defects in this assay when injected separately into two blastomeres at 2-cell stage (18% and 12% CE defects, respectively). In contrast, co-injecting both MOs at subthreshold doses caused strong disruption of CE movements (67%), suggesting that Prickle1 and Ect2 functionally interact during *Xenopus* embryonic development.

**Figure 4:**
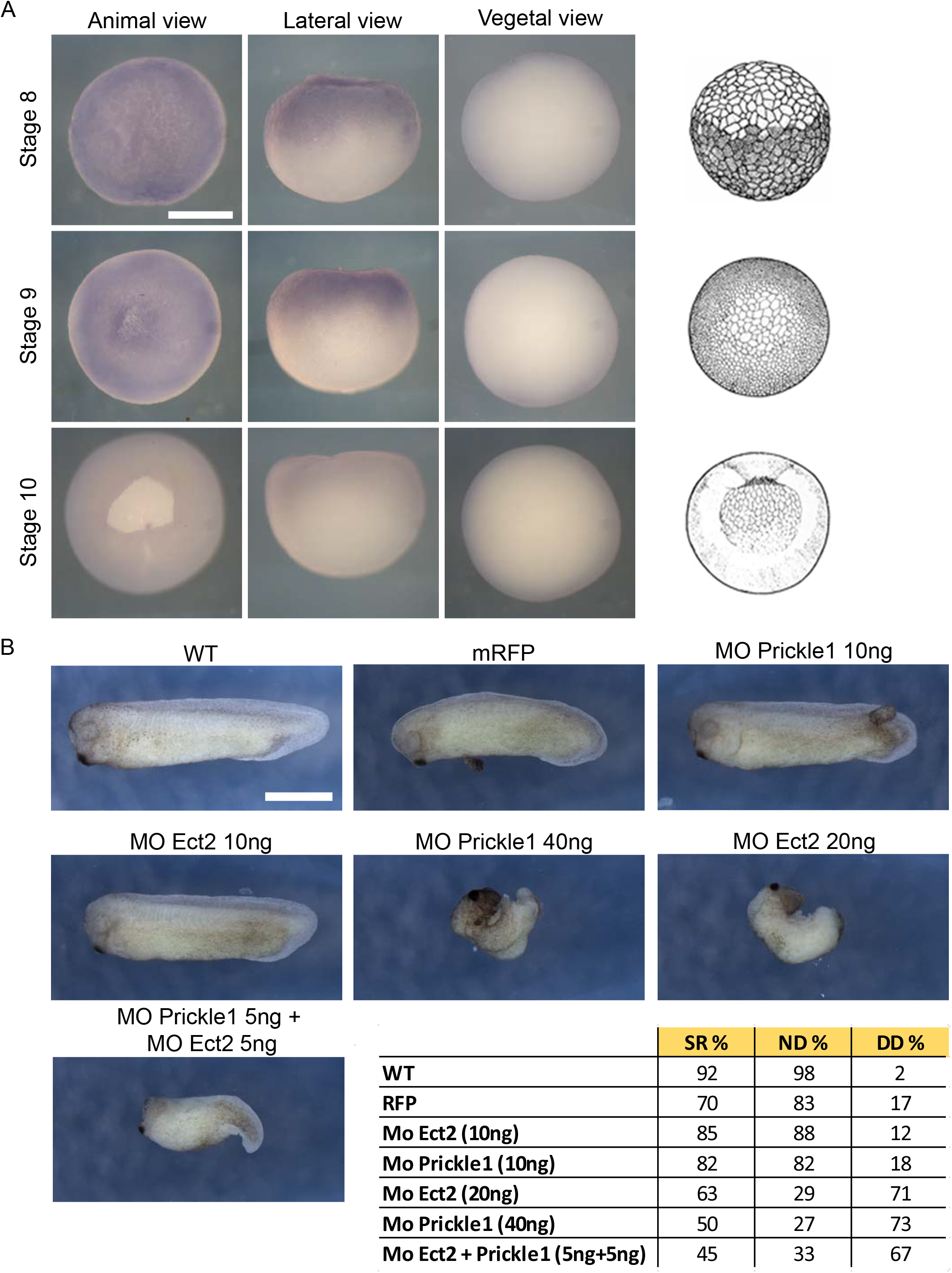
Prickle1 and Ect2 functionally interact in Xenopus during embryonic development. A) *In situ* hybridization against *ect2* transcripts at stage 8, 9 and 10. *ect2* RNA is detectable in the animal pole (animal view and lateral view) but not in the vegetal pole (vegetal view) at stages 8 and 9, but no longer at stage 10. Schematic representations of embryos at the stages analyzed are shown on the right. B) Embryos at 2-cell stage were injected into two blastomeres with Prickle1 and Ect2 MOs as indicated. In all cases 0,5ng of *mRFP* mRNA was injected as control and tracer. Suboptimal doses (10ng) of either MO did not cause CE problems. However, when both Prickle1 and Ect2 MOs were co-injected at suboptimal doses (5ng each), embryos displayed CE problems at a rate comparable to high doses of each MO injected separately (40ng Prickle1-MO or 20ng Ect2-MO). A total of 60 embryos per condition were analyzed in two independent experiments. Pictures illustrate representative phenotypes. (SR=survival rate; ND=percentage of surviving embryos developing normally; CED= percentage of surviving embryos showing convergent-extension defects). Scale bars: A = 0,25 mm; B = 0,5mm.

## Discussion

We and others have demonstrated the prominent role of PRICKLE1 during cancer progression^2, 9–11^. In this study, we identified the protein complex associated to PRICKLE1 and we aimed to evaluate the impact of PRICKLE1 and its associated protein complex in TNBC. Our results show that PRICKLE1 acts as a scaffold protein due to the large number of associated proteins with enzymatic activity. Among the PRICKLE1-associated proteins, we focused our attention on small G-protein regulators since their impact on cell motility and cancer cell dissemination has been well characterized^31–33^. Exploiting our transcriptomic breast cancer database, we showed that this subset of genes is up-regulated in TNBC. Among this group of genes, we identified *ECT2* as the most prominent contributor to *PRICKLE1* prognostic value. Indeed TNBC patient with up-regulated expression of both *PRICKLE1* and *ECT2* expression have a shorter MFS than other patients. We further characterized PRICKLE1 and ECT2 interaction and showed that PRICKLE1 controlled ECT2 function on Rac1 activation. We finally defined that Prickle1 and Ect2 interaction was evolutionary conserved, since both proteins contribute to *Xenopus* embryonic development and are involved in convergent-extension movements.

Among breast cancers, TNBC are considered as the most aggressive form and no targeted therapy is currently available due to a lack of specific targets^1^. Here, we show that *PRICKLE1* is overexpressed in TNBC and is a poor-prognosis marker. PRICKLE1 is a protein highly regulated by post-translational modifications, particularly through ubiquitination/deubiquitination. PRICKLE1 is indeed the target of SMURF1, an ubiquitin ligase, which allows its rapid degradation^34^. PRICKLE1 is also protected from degradation by USP9x which de-ubiquitinates the protein^35^. Interestingly USP9x is also up-regulated in several cancers and is considered as a poor-prognosis marker^36^. PRICKLE1 is also regulated through phosphorylation by the serine/threonine kinase called MINK1, which promotes its function, its membrane localization and association with signaling molecules^12^. Together, this shows that PRICKLE1 is a pivotal protein in cancer cell dissemination and a candidate for setting up novel therapeutic strategies.

During developmental processes and cancer progression, PRICKLE1 is required for oriented cell migration^2, 9, 11, 37^. At the molecular level, we and others have shown that PRICKLE1 contributes to localize VANGL at the plasma membrane^8, 12^, LL5β at the +ends of the microtubules^11^, and to restrict localization of the Rho-GAP at the edge of the migrating cancer cells^10^. PRICKLE1 also regulates spatial localization of several active proteins such as mTORC2 to allow local activation of Akt at the leading edge of migrating cells^2^, PHLDB2 to disassemble focal adhesions^11^ and to restrict RhoA activity by regulating subcellular localization of Rho-GAP^10^. Together the contribution of PRICKLE1 to localization of its interacting partners allows the cells to coordinate cellular movements to create a cellular imbalance and promote directed cell migration. Here we showed that PRICKLE1 also contribute to regulate the activity of ECT2, a GEF for Rac1, which is essential for cell motility.

ECT2 is a Rho-GEF controlling Rac1 activity^13^. Although ECT2 has been extensively studied for its role in the nucleus and during cytokinesis, reports have shown that ECT2 can also be localized in the cytoplasm of cancerous cells^16^. We observed that ECT2 is localized in actin-rich structures within the lamellipodia. As described for other PRICKLE1 interactors, PRICKLE1 might contribute to ECT2 spatial localization in order to modulate its Rac activity. Moreover, our data show that overexpression of ECT2 in HEK293T cells contributes to an increase of Rac activity, and that PRICKLE1 overexpression leads to a decrease of this gain of function, suggesting an inhibitory role of PRICKLE1 on ECT2 activity. Altogether, this depicts PRICKLE1 as a master regulator of localized expression and regulation of signaling events in migratory cancer cells.

Our data also identified a role for the PET domain of PRICKLE1, as ECT2 is to date the only protein identified to be associated with this domain. At the molecular level, it has been shown that PRICKLE1 exists in an open and closed conformation^38^. It has been suggested that in closed conformation, the three LIM domains of PRICKLE1 mask the PRICKLE1 PET domain. In open conformation, the PET domain is unmasked, thus activating PRICKLE1. We can speculate that the interaction between PRICKLE1 and ECT2 can be modulated by switching between these two conformations providing a still uncharacterized molecular mechanism of PRICKLE1 activation.

Finally, our study identified that ECT2 acts during *Xenopus* embryonic development. Prickle1 has been extensively characterized for its contribution during convergent-extension^6, 7^ movements and has been shown to be asymmetrically distributed within cells in order to organize their movement^39, 40^. A previous study indicated that *Prickle1* mRNA accumulates within the blastopore lip from the onset of gastrulation^41^. Here, we showed that *ect2* mRNA and presumably Ect2 protein are expressed prior to and in a broader pattern than Prickle1^41^. Knockdown experiments strongly suggest that Prickle1 and Ect2 act together to allow convergence-extension movements during gastrulation. Altogether, our data support the view that Ect2 might represent a permissive factor for Prickle1 activity. This study demonstrates the importance of the evolutionary conserved interaction between Prickle1 and Ect2, which appears to be reactivated during tumorigenesis to promote cancer cell dissemination and metastasis.

## Supporting information

## List of abbreviations

TNBC: Triple-negative breast cancers
PCP: planar cell polarity
GEF: Guanylyl Exchange Factor
ECT2: Epithelial cell transforming sequence 2
GEF: Guanylyl Exchange Factor
TN: triple-negative
MO: Morpholino antisense oligonucleotides
MFS: metastasis free survival
CE: convergent-extension movements

## Additional Information

### Ethics approval and consent to participate

Not applicable

### Consent for publication

Not applicable

### Availability of data and materials

The mass spectrometry proteomics data, including search results, will be deposited to the ProteomeXchange Consortium (www.proteomexchange.org)^42^ *via* the PRIDE partner repository with the dataset identifier PXD011253.

## Authors’ contributions

AMD, JPB and FB designed the study and wrote the manuscript. AMD conducted the research and performed biochemistry experiments. MSW, LC, SA performed the protein complex purification and mass spectrometry analysis. PF and FB analyzed transcriptomic data base. DR and LK contribute to the Xenopus experiments. DB contributes with key insight.

## Acknowledgements

The authors wish to thank Valérie Ferrier for critical review of the manuscript and Emilie Beaudelet for technical assistance with protein sample for mass spectrometry analysis.

