## Supplementary material for "ECT2 associated to PRICKLE1 are poor-prognosis markers in triple-negative breast cancer"

**Supplementary Table 1: List of breast cancer data sets included in the study**

| Reference | Source of data | N° of samples | Technological platform | N° of probe sets | N° of samples used |
|---|---|---|---|---|---|
| van de Vijver et al., NEJM 2002 | http://microarray-pubs.stanford.edu/wound_NKI/ | 295 | Agilent Hu25K | 25K | 254 |
| van't Veer et al., Nature 2002 | http://www.rii.com/publications/2002/vantveer.html | 117 | Agilent Hu25K | 25K | 117 |
| Expression Project for Oncology (expO), 2005 | https://expo.intgen.org/geo GEO: GSE2109 | 348 | Affymetrix U133 Plus 2.0 | 54K | 348 |
| Farmer P et al., Oncogene 2005 | GEO: GSE1561 | 49 | Affymetrix U133A | 22K | 49 |
| Minn AJ et al., Nature 2005 | GEO: GSE2603 | 99 | Affymetrix U133A | 22K | 99 |
| Wang Y et al., Lancet 2005 | GEO: GSE2034 | 286 | Affymetrix U133A | 22K | 286 |
| Hess KR et al., J Clin Oncol 2006 | MDA133 | 133 | Affymetrix U133A | 22K | 131 |
| Ivshina et al., Cancer Res 2006 | GEO: GSE4922, GSE1456 | 448 | Affymetrix U133 A+B | 2x22K | 448 |
| Sotiriou C et al., J Natl Cancer Inst 2006 | GEO: GSE2990 | 189 | Affymetrix U133A | 22K | 80 |
| Bonnefoi et al., Lancet Oncol 2007 | GEO: GSE6861, GSE4779 | 161 | Affymetrix X3P | 61K | 125 |
| Desmedt C etal., Clin Cancer Res 2007 | GEO: GSE7390 | 198 | Affymetrix U133A | 22K | 154 |
| Miller WR et al., Breast Cancer Res 2010 | GEO: GSE5462 | 116 | Affymetrix U133A | 22K | 116 |
| Klein A et al., Int J Cancer 2007 | GEO: GSE6596 | 26 | Affymetrix U133A | 22K | 24 |
| Bos et al., Nature 2009 | GEO: GSE12276 | 204 | AffymetrixU133 Plus 2.0 | 54K | 204 |
| Hoeflich et al., Clin Cancer Res 2009 | GEO: GSE12763 | 30 | Affymetrix U133 Plus 2.0 | 54K | 30 |
| Marty et al., Breast Cancer Res 2008 | GEO: GSE13787 | 23 | Affymetrix U133 Plus 2.0 | 54K | 23 |
| Merritt WM et al., N Engl J Med 2008 | Array Express: E-MTAB-158 | 130 | Affymetrix U133AAofAv2 | 23K | 130 |
| Schmidt M etal., Cancer Res 2008 | GEO: GSE11121 | 200 | Affymetrix U133A | 22K | 200 |
| Yu K et al., PLoS Genet 2008 | GEO: GSE5364 | 196 | Affymetrix U133A | 22K | 183 |
| Zhang Y et al., Breast Cancer Res Treat 2009 | GEO: GSE12093 | 136 | Affymetrix U133A | 22K | 136 |
| Barry et al., J Clin Oncol 2010 | GEO: GSE23593 | 50 | Affymetrix U133 Plus 2.0 | 54K | 50 |
| Iwamoto T et al., J Natl Cancer Inst 2011 | GEO: GSE22093, GSE22597 | 247 | Affymetrix U133A | 22K | 100 |
| Korde et al., Breast Cancer Res Treat 2010 | GEO: GSE18728 | 61 | Affymetrix U133 Plus 2.0 | 54K | 61 |
| Prat A et al., Breast Cancer Res 2010 | GEO: GSE18229 | 337 | Agilent Hu25K | 25K | 264 |
| Silver et al., J Clin Oncol 2010 | GEO: GSE18864 | 84 | Affymetrix U133 Plus 2.0 | 54K | 84 |
| Tabchy A et al., Clin Cancer Res 2010 | GEO: GSE20271 | 178 | Affymetrix U133A | 22K | 178 |
| Jonsson et al., BCR 2010 | GEO: GSE22133 | 359 | Swegene H_v2.1.1 55K | 55K | 346 |
| Chen et al., Breast Cancer Res Treat 2010 | GEO: GSE10780 | 185 | Affymetrix U133 Plus 2.0 | 54K | 42 |
| Desmedt et al., J Clin Oncol 2011 | GEO: GSE16446 | 120 | Affymetrix U133 Plus 2.0 | 54K | 120 |
| Guedj et al., Oncogene 2011 | Array Express: E-MTAB-365 | 537 | Affymetrix U133 Plus 2.0 | 54K | 452 |
| Hatzis C et al., JAMA 2011 | GEO: GSE25066 | 508 | Affymetrix U133A | 22K | 504 |
| Popovici V et al., Breast Cancer Res 2010 | GEO: GSE20194 | 278 | Affymetrix U133A | 22K | 91 |
| TCGA, Nature 2012 | TCGA Data Portal - BRCA - | 1215 | Illumina, RNAseq V2 | 20K | 1092 |
| Ellis et al., Nature 2012 | GEO: GSE29442, GSE35186 | 201 | Agilent-014850 4x44K | 44K | 201 |
| Curtis et al., Nature 2012 | EGA: EGAS00000000083 | 2136 | Illumina HT 12 | 49K | 1974 |
| Sabatier R et al., (our IPC series) PLoS One 2011 | GEO: GSE31448 | 353 | Affymetrix U133 Plus 2.0 | 54K | 286 |
| TOTAL | | 10233 | | | 8982 |
