## Supplementary material for "ECT2 associated to PRICKLE1 are poor-prognosis markers in triple-negative breast cancer"

**Supplementary material and methods**

**Plasmid constructs and reagents**

The cDNA encoding human PRICKLE1 were obtained from J. L. Wrana. The cherry-ECT2 was obtained from Pr Mark Burkard (Univ. of Wisconsin, USA). Antibodies purchased were mouse anti-FLAG M2 (Sigma), rabbit anti-ECT2 (Merck Millipore) and Anti-Rac1	(Cytoskeleton Inc). Anti-Cherry was a gift from Dr Vincent Géli (CRCM, Marseille, France). The following sequences of siRNA were used to target PRICKLE1: PRICKLE1 (#02) GAGAGAAGCAUCGGAUUAA, PRICKLE1	(#04) GAAGAUAAAUGGAGGUGAA.

**Tissue culture and transfection**

HEK293T and MDA-MB-361 cells were grown in Dulbecco’s modified Eagle medium (DMEM; Sigma) supplemented with 10% fetal bovine serum (FBS; Thermo) and penicillin-streptomycin (Thermo) in a 37°C humidified incubator with 5% CO2. Stable cell lines were generated using lentiviral infection of GFP-Prickle1. Cells lines were obtained from ATCC and are regularly tested for mycoplasma. Cells are used for no longer passage than 20.

**Immunopurification**

5x10^6^ MDA-MB-231 cells expressing either GFP-PRICKLE1 were used for affinity purification procedure. Briefly, cells were lysed and solubilized in TAP lysis buffer (0.1% Igepal CA 630, 10% glycerol, 50mM Hepes-NaOH; pH 8.0, 150mM NaCl, 2mM EDTA, 2mM DTT, 10mM NaF, 0.25mM NaOVO3, 50mM β-glycerophosphate, and protease inhibitor cocktail (Calbiochem). After 30 min centrifugation at 40,000xg (18,000 rpm in a Beckman JA20 rotor), the soluble fraction was incubated overnight at 4°C with anti-GFP nanobodies (Gift from Dr Mauro Modesti, CRCM, Marseille, France). Beads were washed with 100 volumes of TAP lysis buffer. Beads elution was performed using laemmli buffer.

**Affinity purification, immunoprecipitation and western blot**

48 hours post-transfection, cells were lysed with the TAP lysis buffer and incubated at 4ºC for 1 hour to solubilize proteins. Affinity purification and immunoprecipitations were performed using anti-FLAG-M2 beads (Sigma) for 3 hours at 4°C. After extensive washes with lysis buffer, proteins were eluted with 2x Laemmli sample buffer and heated at 95ºC for 5 min in the presence of β-mercaptoethanol (Sigma). Whole cell lysates or purified protein samples were resolved by SDS-polyacrylamide gel electrophoresis (SDS-PAGE) and transferred onto Biotrace NT Nitrocellulose Transfer Membranes (GE Healthcare). Western blotting was performed with antibodies as indicated in figures legend, followed by chemiluminescent detection using appropriate HRP-conjugated antibodies and the ECL (GE Healthcare) reagent.

**Mass spectrometry analysis.**

Protein extract were loaded on NuPAGE 4-12% Bis-Tris acrylamide gels (Life Technologies) to stack proteins in a single band that was stained with Imperial Blue (Pierce, Rockford, IL) and cut from the gel. Gels pieces were submitted to an in-gel trypsin digestion(Shevchenko et al 1996) with slight modifications. Briefly, gel pieces were washed and destained using 100 mM NH_4_HCO_3_. Destained gel pieces were shrunk with 100 mM ammonium bicarbonate in 50% acetonitrile and dried at room temperature. Protein spots were then rehydrated using 10 mM DTT in 100 mM ammonium bicarbonate pH 8.0 for 45 min at 56 °C. This solution was replaced by 55 mM iodoacetamide in 100 mM ammonium bicarbonate pH 8.0 and the gel pieces were incubated for 30 min at room temperature in the dark. They were then washed twice in 100 mM ammonium bicarbonate and finally shrunk by incubation for 5 min with 100 mM ammonium bicarbonate in 50% acetonitrile. The resulting alkylated gel pieces were dried at room temperature. The dried gel pieces were re-swollen by incubation in 100 mM ammonium bicarbonate pH 8.0 supplemented with trypsin (12.5 ng/µL; Promega) for 1 h at 4°C and then incubated overnight at 37 C. Peptides were harvested by collecting the initial digestion solution and carrying out two extractions; first in 5% formic acid and then in 5% formic acid in 60% acetonitrile. Pooled extracts were dried down in a centrifugal vacuum system. Samples were reconstituted with 0.1% trifluoroacetic acid in 4% acetonitrile and analyzed by liquid chromatography (LC)-tandem mass spectrometry (MS/MS) using an Orbitrap Fusion Lumos Tribrid Mass Spectrometer (Thermo Electron, Bremen, Germany) online with a an Ultimate 3000RSLCnano chromatography system (Thermo Fisher Scientific, Sunnyvale, CA). Peptides were separated on a Dionex Acclaim PepMap RSLC C18 column. First peptides were concentrated and purified on a pre-column from Dionex (C18 PepMap100, 2 cm × 100 µm I.D, 100 Å pore size, 5 µm particle size) in solvent A (0.1% formic acid in water). In the second step, peptides were separated on a reverse phase LC EASY-Spray C18 column from Dionex (PepMap RSLC C18, 15 cm × 75 µm I.D, 100 Å pore size, 2 µm particle size) at 300 nL/min flow rate. After column equilibration using 4% of solvent B (20% water - 80% acetonitrile - 0.1% formic acid), peptides were eluted from the analytical column by a two steps linear gradient (4-22% acetonitrile/H_2_O; 0.1 % formic acid for 110 min and 22-32% acetonitrile/H_2_O; 0.1 % formic acid for 10 min). For peptide ionization in the EASY-Spray nanosource, spray voltage was set at 2.2 kV and the capillary temperature at 275 °C. The Orbitrap Lumos was used in data dependent mode to switch consistently between MS and MS/MS. Time between Masters Scans was set to 3 seconds. MS spectra were acquired with the Orbitrap in the range of m/z 400-1600 at a FWHM resolution of 120 000 measured at 400 *m/z*. AGC target was set at 4.0e5 with a 50 ms Maximum Injection Time. For internal mass calibration the 445.120025 ions was used as lock mass. The more abundant precursor ions were selected and collision-induced dissociation fragmentation was performed in the ion trap to have maximum sensitivity and yield a maximum amount of MS/MS data. Number of precursor ions was automatically defined along run in 3s windows using the “Inject Ions for All Available pararallelizable time option” with a maximum injection time of 300 ms. The signal threshold for an MS/MS event was set to 5000 counts. Charge state screening was enabled to exclude precursors with 0 and 1 charge states. Dynamic exclusion was enabled with a repeat count of 1 and a duration of 60 s.

**Protein identification**

*Data Analysis-*Raw files generated from mass spectrometry analysis were processed with Proteome Discoverer 1.4.1.14 (Thermofisher Scientific). This software was used to search data via in-house Mascot server (version 2.4; Matrix Science Inc., London, UK) against the Human database subset (20244 sequences) of the SwissProt database (version 2017.11). Database search were done using the following settings: a maximum of two trypsin miscleavage allowed, methionine oxidation (+15.99491), N-terminal acetylation (+42.0106) as variable modifications, cysteine carbamido-methylation (+57.02146) as a fixed modification. A peptide mass tolerance of 6 ppm and a fragment mass tolerance of 0.8 Da were allowed for search analysis. Only peptides with higher Mascot threshold (identity) were selected. Only proteins with a FDR < 1% were selected for identification. Relative intensity-based label-free quantification (LFQ) was processed using the MaxLFQ algorithm(Cox et al 2014) from the freely available MaxQuant computational proteomics platform, version 1.6.2.1(Cox and Mann 2008). The acquired raw LC Orbitrap MS data were first processed using the integrated Andromeda search engine(Cox et al 2011). Spectra were searched against the Human database	(UniProt Proteome reference, date 2018.09; 20395 entries). This database was supplemented with a set of 245 frequently observed contaminants. The following parameters were used for searches: (*i*) trypsin allowing cleavage before proline; (ii) two missed cleavages were allowed; (*ii*) monoisotopic precursor tolerance of 20 ppm in the first search used for recalibration, followed by 4.5 ppm for the main search and 0.5 Da for fragment ions from MS/MS ; (*iii*) cysteine carbamidomethylation	(+57.02146) as a fixed modification and methionine oxidation (+15.99491) and N-terminal acetylation (+42.0106) as variable modifications; (*iv*) a maximum of five modifications per peptide allowed; and (*v*) minimum peptide length was 7 amino acids and a maximum mass of 4,600 Da. The match between runs option was enabled to transfer identifications across different LC-MS/MS replicates based on their masses and retention time within a match time window of 0.7 min and using an alignment time window of 20min. The quantification was performed using a minimum ratio count of 1 (unique+razor) and the second peptide option to allow identification of two co-fragmented co-eluting peptides with similar masses. The false discovery rate (FDR) at the peptide and protein levels were set to 1% and determined by searching a reverse database. For protein grouping, all proteins that cannot be distinguished based on their identified peptides were assembled into a single entry according to the MaxQuant rules. The statistical analysis was done with Perseus program (version 1.6.1.3) from the MaxQuant environment ([www.maxquant.org](http://www.maxquant.org)). The LFQ normalised intensities were uploaded from the proteinGroups.txt file. First, proteins marked as contaminant, reverse hits, and “only identified by site” were discarded. Quantifiable proteins were defined as those detected in at least 100% of samples in at least one condition. Protein LFQ normalized intensities were base 2 logarithmized to obtain a normal distribution. Missing values were replaced using data imputation by randomly selecting from a normal distribution centred on the lower edge of the intensity values that simulates signals of low abundant proteins using default parameters (a downshift of 1.8 standard deviation and a width of 0.3 of the original distribution). To determine whether a given detected protein was specifically differential a two-sample t-test were done using permutation based FDR-controlled at 0.001 and employing 250 permutations. The p value was adjusted using a scaling factor s0 with a value of 1 (Tusher et al 2001) . The mass spectrometry proteomics data, including search results, will be deposited to the ProteomeXchange Consortium ([www.proteomexchange.org](http://www.proteomexchange.org))(Vizcaino et al 2014) *via* the PRIDE partner repository with the dataset identifier PXD011253.

***In situ* hybridization (ISH)**

The preparation of digoxigenin-labeled antisense RNA probes and the whole-mount ISH were performed as previously described(Pizard et al 2004), except that the proteinase K step was omitted. Whole-mount chromogenic *in situ* hybridization was performed as detailed by Marchal et al., 2009. Probes were generated from linearized plasmids using RNA-labeling mix (Roche). 40ng of *ect2* digoxigenin-labeled sense and antisense riboprobes were used for hybridization. Anti-Dig-AP (Roche, 11266026, 1:5000) antibody was used to stain embryos, which were then fixed, washed in methanol, rehydrated and photographed.
